# Alpha9alpha10 knockout mice show altered physiological and behavioral responses to signals in masking noise

**DOI:** 10.1101/2023.11.21.567909

**Authors:** Jane A. Mondul, Kali Burke, Barbara Morley, Amanda M. Lauer

## Abstract

Medial olivocochlear (MOC) efferents modulate outer hair cell motility through specialized nicotinic acetylcholine receptors to support encoding of signals in noise. Transgenic mice lacking the alpha9 subunits of these receptors (α9KOs) have normal hearing in quiet and noise, but lack classic cochlear suppression effects and show abnormal temporal, spectral, and spatial processing. Mice deficient for both the alpha9 and alpha10 receptor subunits (α9α10KOs) may exhibit more severe MOC-related phenotypes. Like α9KOs, α9α10KOs have normal auditory brainstem response (ABR) thresholds and weak MOC reflexes. Here, we further characterized auditory function in α9α10KO mice. Wildtype and α9α10KO mice had similar ABR thresholds and acoustic startle response (ASR) amplitudes in quiet and noise, and similar frequency and intensity difference sensitivity. α9α10KO mice had larger ABR Wave I amplitudes than wildtypes in quiet and noise, but the noise:quiet amplitude ratio suggested α9α10KOs were more susceptible to masking effects for some stimuli. α9α10KO mice also had larger startle amplitudes in tone backgrounds than wildtypes. Overall, α9α10KO mice had grossly normal auditory function in quiet and noise, though their larger ABR amplitudes and hyperreactive startles suggest some auditory processing abnormalities. These findings contribute to the growing literature showing mixed effects of MOC dysfunction on hearing.

## I. INTRODUCTION

The medial olivocochlear (MOC) efferent system modulates afferent encoding of incoming sounds primarily via cholinergic inhibition of the outer hair cells (Fuchs & Lauer, 2019). MOC neurons reduce outer hair cell motility through specialized nicotinic acetylcholine receptors (nAChRs) comprised of alpha9 and alpha10 subunits (Elgoyhen et al., 1994; Elgoyhen et al., 2001), thus serving as a gain control for the cochlear amplifier. Previous work using lesions or stimulations of the olivocochlear bundle have indicated a role in encoding of signals in noise and protection from acoustic injury (Boero et al., 2018; Elgoyhen, 2020; Guinan, 2006; Kawase & Liberman, 1993; Liberman & Guinan, 1998; Lopez-Poveda, 2018; Maison et al., 2013). However, demonstrating clear and consistent behavioral effects of olivocochlear manipulations has been less straightforward (Lauer et al., 2022).

Much of what is known about MOC function has been derived from studies in alpha9 knockout (α9KO) transgenic mice (Vetter et al., 1999). These mice were generated to have a null mutation in the alpha9 subunit of nAChRs that results in the loss the classic inhibitory effects on cochlear activity as measured via suppression of compound action potentials and otoacoustic emissions by electrical stimulation. Binding of acetylcholine to the outer hair cell nAChRs reduces calcium influx, thereby hyperpolarizing the cell (Fuchs, 2014). α9KO mice show structural changes to MOC innervation patterns, including fewer but larger MOC terminals per outer hair cell and disrupted patterns of tunneling fibers (Boero et al., 2018; Lauer & May, 2011; Morley et al., 2017; Vetter et al., 1999), suggesting that functional alpha9 nAChR subunits are necessary for normal development and function of the MOC system.

The hearing abilities of α9KOs have been probed using a variety of physiological and behavioral methods in order to better understand the role of the MOC system in hearing. α9KO mice have normal hearing sensitivity for tones in quiet as measured by auditory brainstem responses (ABRs; Lauer, 2017; Lauer & May, 2011; May et al., 2002; Morley et al., 2017; Vetter et al., 1999), distortion product otoacoustic emissions (DPOAEs; Lauer & May, 2011; Morley et al., 2017), and behavior (May et al., 2002; Prosen et al., 2000). Surprisingly, α9KOs also show normal tone detection, intensity discrimination, and prepulse inhibition in noise (Allen & Luebke, 2017; May et al., 2002). However, α9KOs show weak MOC reflexes (Chambers et al., 2012; Vetter et al., 1999; but see Morley et al., 2017), abnormal temporal processing (Lauer & May, 2011), reduced prepulse inhibition in quiet (Allen & Luebke, 2017), and impaired frequency resolution and spatial hearing (Clause et al., 2017). Overall, α9KO mouse studies seem to refute the hypothesis that the MOC system plays a role in hearing-in-noise or suggest that other compensatory mechanisms take over when MOC activity is disrupted. The deficits observed in other suprathreshold processing abilities are likely related to abnormal development of the central auditory system (Clause et al., 2014).

Another transgenic model that is deficient for both the alpha9 and alpha10 subunits of nAChRs was generated by Morley et al. (2017). The alpha9alpha10 double knockout model (α9α10KO) was developed to investigate whether the deletion of both subunits would produce a more severe MOC lesion phenotype than the α9KO. Morley et al. (2017) found that α9α10KOs have comparable ABR thresholds and DPOAE amplitudes to both wildtypes and α9KOs. However, α9α10KOs have weaker MOC reflexes than wildtypes, α9KOs, and α10KOs. Further characterization of hearing abilities in the α9α10KO mouse model could provide additional evidence toward hypothesized roles of the MOC system. In the present study, we aimed to characterize auditory function more comprehensively in adult α9α10KO mice by using both physiological and reflex-based behavioral methods to probe hearing-in-noise and suprathreshold processing abilities in untrained animals. Specifically, we measured: 1) ABR thresholds, amplitudes, and latencies to sounds in quiet and in noise, 2) acoustic startle responses (ASRs) to sounds in quiet and in noise, and 3) frequency and intensity difference sensitivity using prepulse inhibition (PPI) of the ASR.

## II. METHODS

### A. Subjects

Experiments were conducted in 3-month-old mice of two strains: C57BL/6J (“WT”; wildtype controls; *n* = 7, 3 female) and alpha9alpha10 double knockouts (“α9α10KO”; *n* = 9, 4 female). Mice were obtained from the Morley laboratory at Boys Town National Research Hospital. WT controls were from the same colony as the α9α10KO mice and obtained by mating heterozygote females with heterozygote or wildtype males or wildtype females with heterozygote males. Background strain, generation, and validation of the transgenic line was previously described (Morley et al., 2017). Briefly, the alpha9 and alpha10 knockout strains were backcrossed to >99% congenicity on the C57BL/6J background using MAXBAX (Charles River, Troy NY). The alpha9 and alpha10 knockout strains were crossed to construct the double knockout. The double knockout strain was uncrossed by breeding with wildtype C57BL/6J mice obtained from Jackson Labs. Knockouts were uncrossed and re-crossed yearly. This results in a background in the double KO mouse with greater similarity than would be found with a simple cross of the alpha9 and alpha10 strains.

Prior to testing, animals were group housed in high-traffic mouse vivaria until 2 months of age, transported to Johns Hopkins University, and acclimated to the new housing facility for four weeks. Mice were exposed to unknown levels of noise during transport, but this was a common factor for all animals tested in this study. At the time of testing, animals were group housed in a low noise vivarium; sound levels were previously described (Wu et al., 2020). The housing room was maintained on a 12:12 hour light:dark cycle (7:00 to 19:00). Up to five mice were housed per one filter top shoebox cage (30 x 19 x 13 cm) with corncob bedding and nestlets. Exclusion criteria included abnormal hearing thresholds in quiet or signs of outer or middle ear infections; however, no animals were excluded from any of these experiments. Mice weighed between 20-30 grams at the time of testing, with no significant difference across strains. All procedures were approved by the Johns Hopkins University Animal Care and Use Committee (ACUC) and follow the NIH ARRIVE Guidelines.

### B. Procedures

#### 1. Auditory brainstem responses (ABRs) in quiet and in masking noise

ABR testing procedures were similar to those previously described in this laboratory (e.g. Capshaw et al., 2022; Vicencio-Jimenez et al., 2021). Briefly, ABR testing was conducted in a sound-treated booth (Industrial Acoustics Company (IAC), Bronx, NY; 59” x 74” x 60”) lined with acoustic foam (Pinta Acoustic, Minneapolis, MN). Mice were anesthetized with an intraperitoneal injection of 100 mg/kg ketamine and 20 mg/kg xylazine and placed on a heating pad to maintain a temperature of 37°C. Subdermal needle electrodes (Disposable Horizon, 13 mm needle, Rochester Med, Coral Springs, FL) were placed on the vertex (active), ipsilateral mastoid (reference), and hind limb (ground) in a standard ABR recording montage. One ear was tested per subject (random and counter-balanced selection). ABR signals were acquired with a Medusa4Z preamplifier (12 kHz sampling rate) and filtered from 300-3000 Hz with an additional band-reject filter at 60 Hz. Posthoc filters from 300-3000 Hz with steeper cutoff slopes were also applied for additional smoothing.

ABRs in quiet were recorded to clicks (100μs) and tonebursts (4, 8, 12, 16, 24, 32 kHz; 5 ms duration, 0.5 ms rise/fall) at a rate of 21/s for a total of 512 presentations with alternating stimulus polarities. Stimuli were created in SigGen software (Tucker-Davis Technologies (TDT), Alachua, FL) and generated by a RZ6 multi-I/O processor (TDT). Stimuli were played from a free field speaker (MF1, TDT) located 10 cm from the animal’s ear canal at 0° azimuth. Stimuli were calibrated with a 0.25” free-field microphone (PCB Piezotronics, Depew, NY Model number 378C01) placed at the location of the animal’s ear canal. Stimulus level ranged from 90 to 10 dB SPL in 10 dB steps.

Masked ABRs were recorded to clicks and tonebursts (4, 8, 12, 16 kHz) in the presence of a 50 dB SPL broadband noise (4-20 kHz; 8 dB SPL spectrum level). Masking noise was generated by an Elgenco 602A gaussian noise generator and presented from a separate free field speaker (MF1, TDT) located adjacent to the stimulus speaker and within the minimum audible angle of mice (Behrens & Klump, 2016; Heffner et al., 2001; Lauer et al., 2011). The sound level was calibrated with a Larson Davis sound level meter (Model 824) with Z-weighting prior to each recording session. The testing order of ABRs in quiet and in noise was randomized across animals.

ABR traces were analyzed offline by two researchers, one of whom was blinded to the subject and stimulus condition. Inter-rater reliability was >0.85 for ABR thresholds, amplitudes, and latencies. ABR threshold was defined as the average between the lowest sound level to evoke a response and the first level with no response (any wave). Peak-to-trough amplitudes and peak latencies were derived for ABR Waves I, II III, and IV using manual peak-picking methods and a semi-automated ABR wave analysis software previously described (Burke et al., 2023). Due to variability in central wave morphologies, amplitude and latency analyses were focused on Wave I only. ABR threshold differences between noise and quiet (dB masking) and ratios between ABR Wave I amplitudes in quiet and in noise were calculated to probe the effects of masking noise.

#### 2. Acoustic startle response (ASR) and prepulse inhibition (PPI) general procedures

ASR and PPI testing procedures were similar to those previously described (e.g. Clause et al., 2017; Kim et al., 2022). Briefly, mice were brought into the testing room to acclimate 30 minutes prior to testing. Animals were tested one at a time, and the order of tasks (ASR in quiet, ASR in noise, PPI frequency, PPI intensity) was pseudorandomized. All startle experiments were conducted by the same experimenter during the daytime light cycle of the animals’ housing, between 8:00 and 18:00.

Testing was conducted in a sound-treated booth (IAC; 37 x 53 x 33 cm) lined with acoustic foam (Pinta Acoustic, Minneapolis, MN). Stimuli and masking noise were delivered from two adjacent speakers (RadioShack Super Tweeter) located on one end of the sound booth. Stimuli were generated by an RP2.1 real time processor (TDT), a PA5 programmable sound attenuator (TDT), and an amplifier (Crown D75A). Speakers were located 15 cm from the animal’s head and intensity was calibrated using a Larson Davis Sound Level Meter (Model 824) with z-weighting.

During testing, mice were placed into an acoustically transparent mouse restraint device (7.2 x 3.3 x 2.8 cm) atop a piezoelectric accelerometer in the center of the sound-treated booth. Movements of the animal were recorded as voltage. For all tests, a random inter-trial waiting period of 5-15 seconds was used to prevent subjects from predicting the onset of a trial, followed by a 5 second quiet period with a noise criterion of <0.4V to ensure the animal was still prior to the onset of the stimulus. The subject’s startle response was recorded over 120 ms following the onset of the startle eliciting stimulus. ASR amplitude was defined as the maximum peak-to-peak voltage during the 120 ms recording window. Testing was conducted over two sessions per animal lasting 35-45 minutes each. Animals were returned to their home cage after testing. Data were screened offline to ensure that all trials counted as startles had appropriate latency and amplitude values.

#### 3. ASR in quiet and in masking noise

ASRs were measured in response to noise bursts (20 ms) in quiet and in the presence of continuous 60 dB SPL broadband noise (4-20 kHz). Pulses were pseudorandomly presented at levels of 70, 80, 90, 100, and 105 dB SPL for a minimum of 10 times per level.

#### 4. ASR frequency and intensity difference sensitivity

ASR tasks probing frequency and intensity difference sensitivity were conducted in the presence of a 65 dB SPL 10 kHz tone background. Startle eliciting stimuli for both tasks were 20 ms broadband noise burst presented at 105 dB SPL. The startle noise bursts were presented immediately after a prepulse cue. For the frequency difference task (FD), the background tone frequency changed from 10 kHz to one of eight off frequencies (7, 8, 9, 9.5, 10.5, 11, 12, 13 kHz) in a pseudorandom order. For the intensity difference task (ID), the background tone level changed from 70 dB SPL to one of five other intensities (72, 74, 76, 78, 80) in a pseudorandom order. Since d’-like estimates of sensitivity do not exceed 1.0 in PPI tests, we did not calculate frequency or intensity difference limens or thresholds for this experiment (Kim et al., 2022; Lauer et al., 2017).

### C. Statistical analyses

To determine whether α9α10KO mice had different hearing sensitivity in quiet or noise conditions, we used linear mixed-effects models (lmer in the lme4 R package, RRID: SCR_015654) to assess the effects of strain, sex, and stimulus frequency or level on dependent variables such as ABR thresholds, ABR Wave I amplitudes and amplitude ratios, ASR magnitude and PPI. Strain, sex, stimulus frequency, and stimulus level were treated as categorical factors. We controlled for individual dependencies in our data by including a random intercept for mouse identity. Model selection was done using the step-up method for linear mixed effects modeling and the goodness of fit for our model was measured using the Akaike information criterion (AICc). We present the results for the ANOVA based on each model, as well as post hoc tests controlling for multiple comparisons using the mvt adjustment (emmeans R package, RRID: SCR_018734). Tukey’s post hoc analyses were performed to assess significance (emmeans R package, RRID: SCR_018734).

## III. RESULTS

### A. Auditory brainstem responses in quiet and in noise

Auditory brainstem responses (ABRs) were measured to clicks and tonebursts in quiet and in 50 dB SPL masking noise. Mean waveforms are shown for each strain in response to a 90 dB SPL click in quiet and in noise (Fig 1A, top and bottom panels). Wave morphology was generally as expected for α9α10KO mice, with all waves present in quiet and in noise but a less robust waveform in noise. α9α10KO mouse ABR waveforms appeared more variable across individual animals than for the WT mice, especially in quiet and for later wave components. For both strains, the addition of masking noise caused an elevation of ABR thresholds (Fig 1B) and reduction of ABR amplitudes (Fig 1C), but had minimal effects on Wave I latency (Fig 1D).

**Figure 1.**
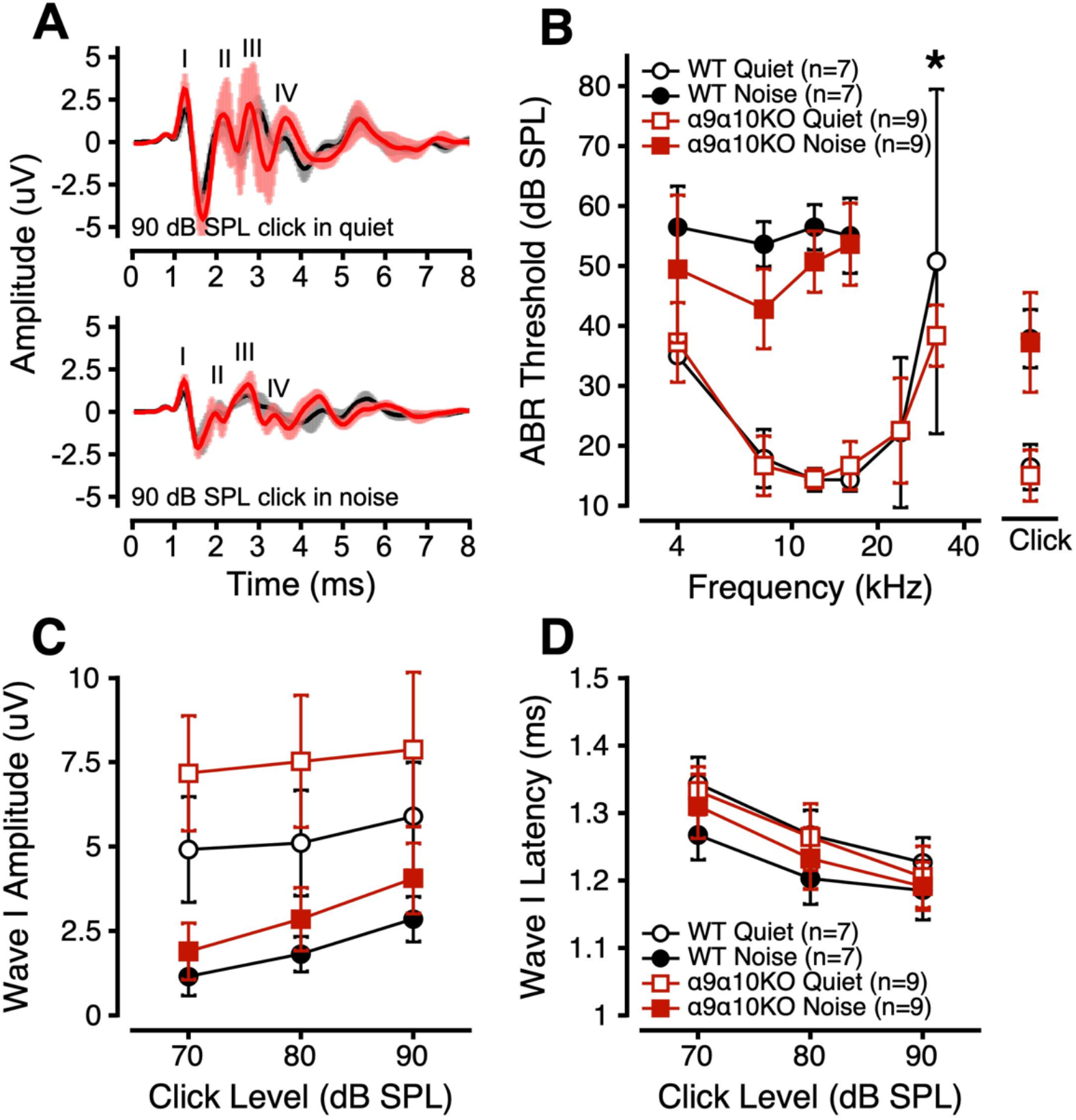
Auditory brainstem responses in quiet and noise for α9α10KO and WT mice. A. Mean (± 1 standard deviation) ABR waveforms in response to a 90 dB SPL click in quiet (top) and noise (bottom) for α9α10KO (trace in foreground) and WT mice (trace in background). B. ABR thresholds as a function of stimulus frequency (B), Wave I amplitudes to suprathreshold clicks (C), and Wave I latencies to suprathreshold clicks (D) in quiet (open symbols) and noise (filled symbols) for α9α10KO (squares) and WT mice (circles). Error bars depict ± 1 standard deviation from the mean.

Overall, ABR thresholds in quiet and in noise were not different between WT and α9α10KO mice, except for higher and more variable thresholds at 32kHz in the WTs (Fig 1B). Linear mixed-effects models comparing ABR thresholds in quiet across strain, stimulus, and sex revealed significant main effects of stimulus and sex, with significant interactions between sex/stimulus and strain/stimulus (see Table I). Post hoc analyses revealed a significant threshold difference at 32kHz across strains (*p* = 0.0033). Linear mixed-effects models comparing ABR thresholds in noise across strain, stimulus, and sex revealed significant main effects of strain and stimulus, with a significant three-way interaction between strain, sex, and stimulus (see Table I). Post hoc analyses revealed a significant difference between male WT and α9α10KO ABR thresholds in noise at 4 and 8kHz (*p* = 0.0095 and 0.0038, respectively).

**TABLE I.**
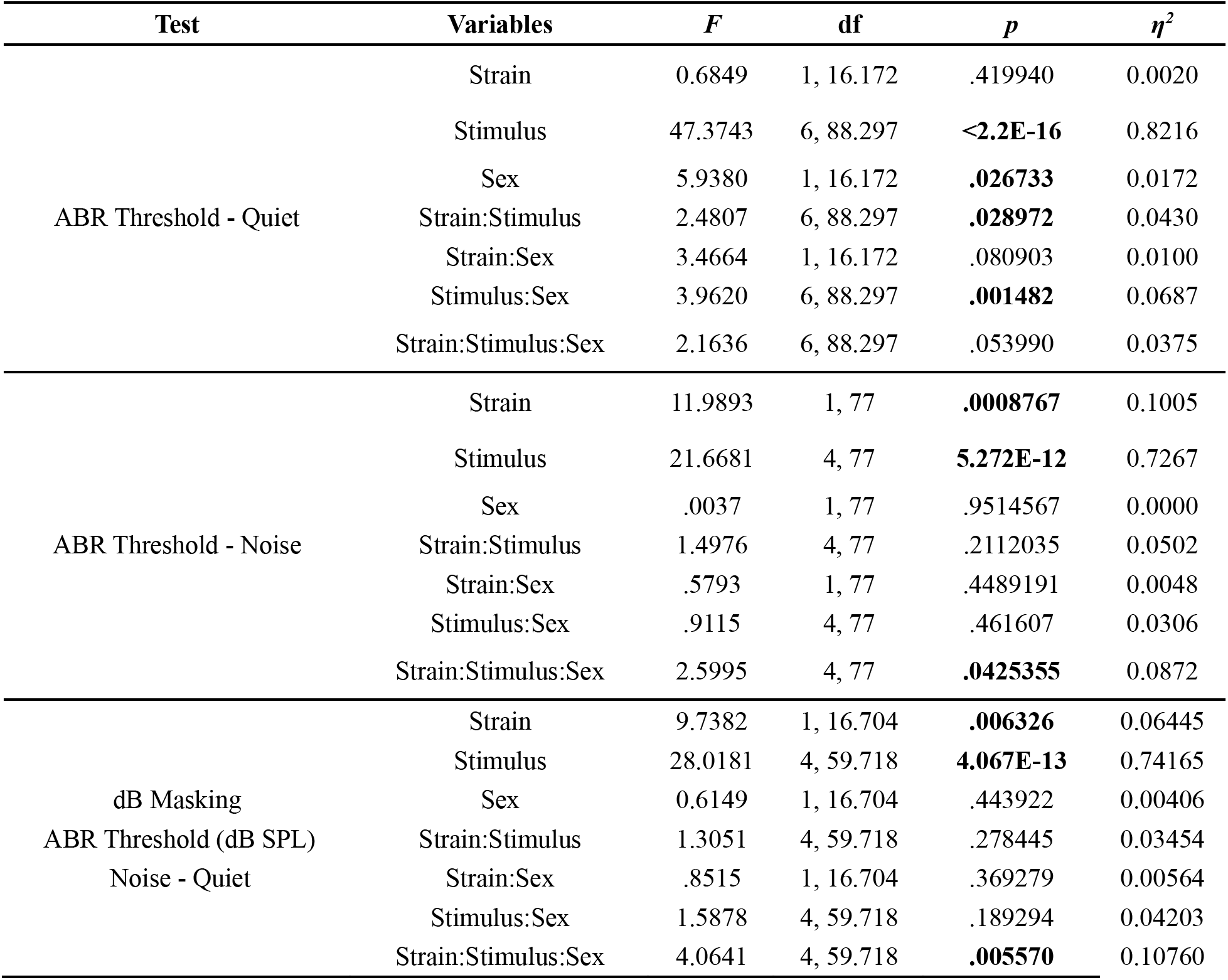
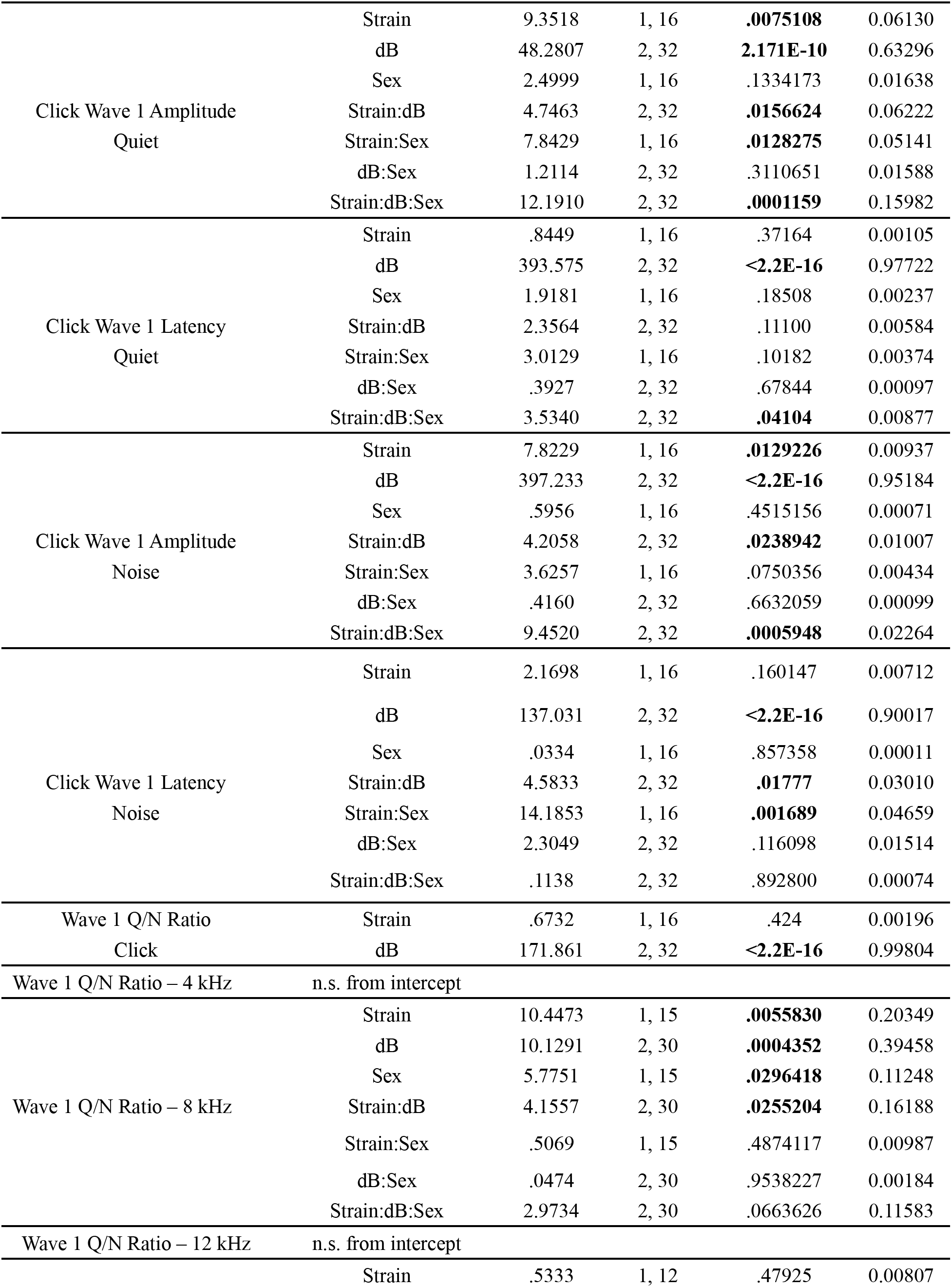
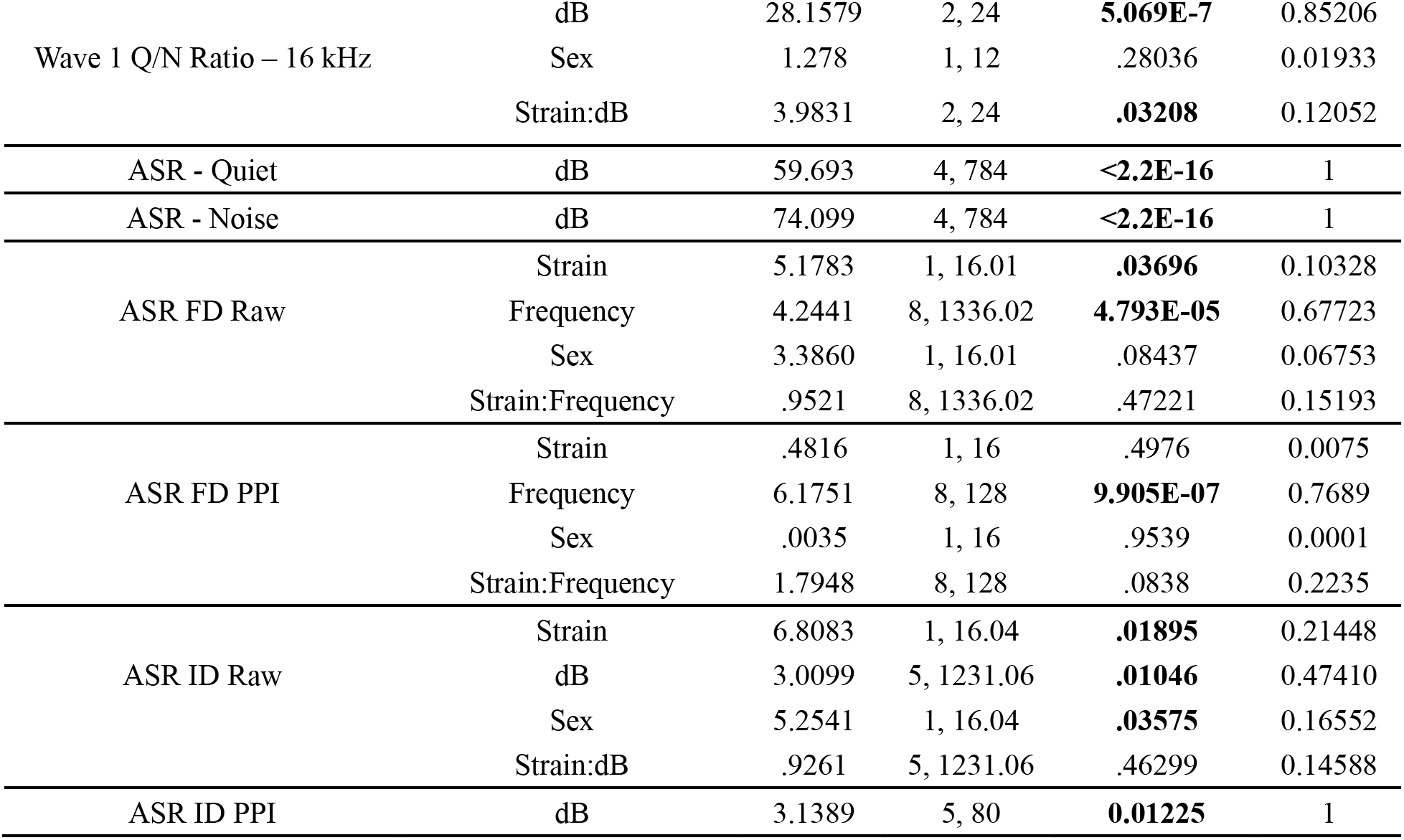
Statistical analysis results.

To evaluate the effects of masking noise on α9α10KO ABR thresholds, we calculated the amount of masking threshold shift for each subject and stimulus (*dB Masking* = *ABR Threshold_Noise_ – ABR Threshold_Quiet_*). Since the addition of masking noise elevates ABR thresholds, dB masking values across strains and stimuli were greater than 0 dB SPL. α9α10KO mice exhibited less masking (i.e. smaller dB masking values) than WTs (Table II). Linear mixed-effects models comparing dB masking values across strain, stimulus, and sex revealed significant main effects of strain and stimulus, with a significant three-way interaction of strain, stimulus, and sex (see Table I). Post hoc analyses indicated significant strain differences in dB masking values for males at 4kHz (*p* = 0.0038) and 8kHz (*p* = 0.0050) and for females at 16kHz (*p* = 0.0187). Overall, these results suggest that α9α10KO mouse ABRs were less susceptible to masking noise than WTs.

**TABLE II.**
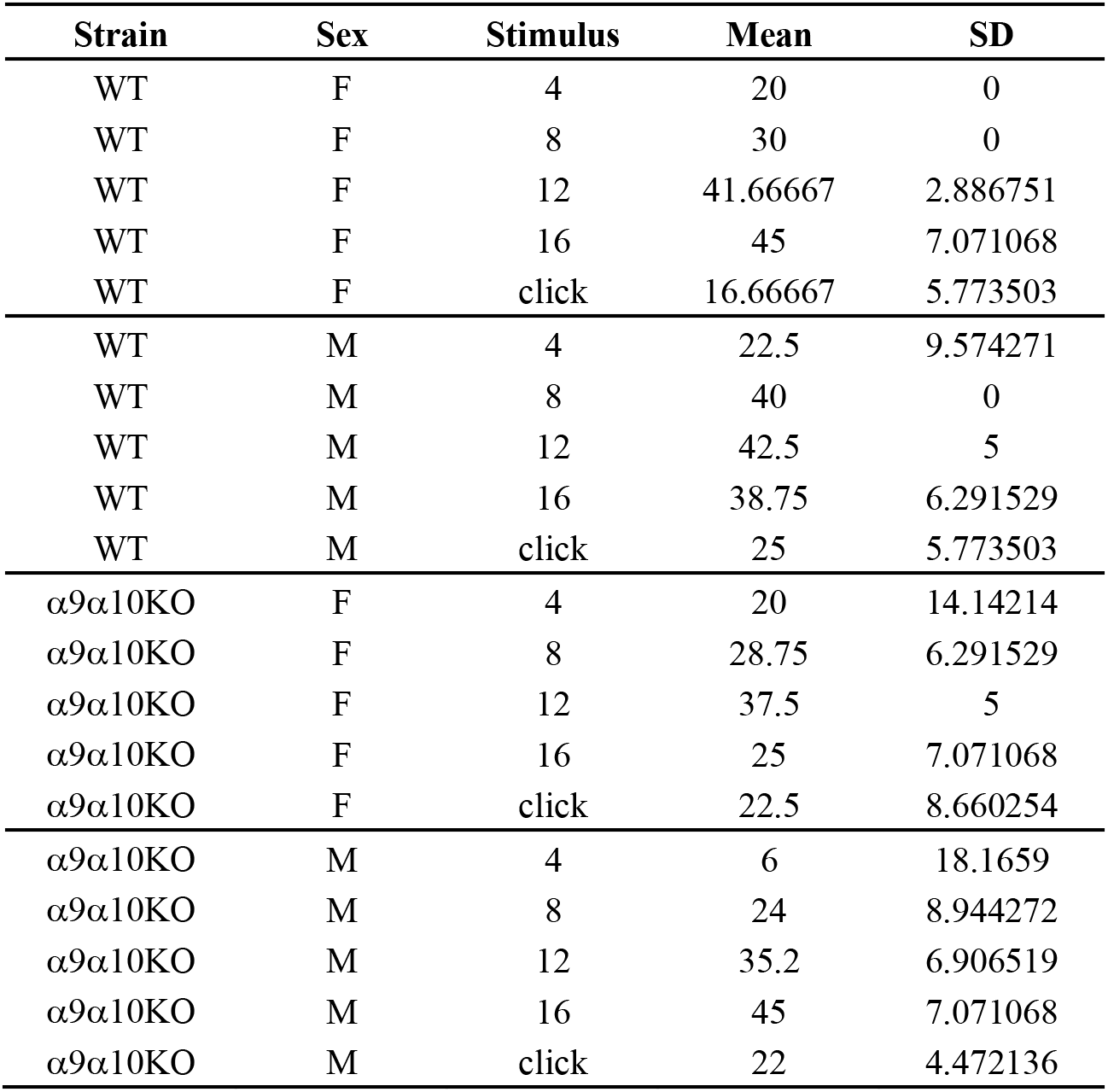
Descriptive statistics of ABR masking (dB SPL) by strain, sex, and stimulus.

ABR Wave I amplitudes and latencies to 90, 80, and 70 dB SPL clicks were quantified for both strains. α9α10KO mice tended to have larger Wave I amplitudes in both quiet and in noise compared to WTs (Fig 1C) and minimal differences in Wave I latency (Fig 1D). For clicks in quiet, linear mixed-effects models comparing ABR Wave I amplitudes across strain, sex, and click level revealed significant main effects of strain and click level, and significant interactions between strain/sex, strain/click level, and strain/sex/click level (see Table I). Post hoc analyses revealed a significant difference in quiet Wave I amplitudes between male WTs and male α9α10KOs for 70, 80, and 90 dB SPL clicks (*p* = 0.0018, 0.0008, 0.0006 respectively). Linear mixed-effects models comparing ABR Wave I latencies across strain, sex, and click level revealed a significant main effect of click level and a significant three-way interaction. Post hoc analyses identified a significant difference in Wave I latency between female WTs and female α9α10KOs for 90 dB SPL clicks only (*p* = 0.475).

For clicks in noise, linear mixed-effects models comparing ABR Wave I amplitudes across strain, sex, and click level revealed significant main effects of strain and click level, and significant interactions between strain/click level and strain/sex/click level (see Table I). Post hoc analyses revealed a significant difference in masked Wave I amplitudes between male WTs and male α9α10KOs for 80 and 90 dB SPL clicks only (*p* = 0.0063, 0.0006 respectively). Linear mixed-effects models comparing ABR Wave I latencies across strain, sex, and click level revealed a significant main effect of click level, and significant interactions between strain/sex and strain/click level. Post hoc analyses of ABR Wave I latencies identified a significant strain difference for males (*p* = 0.0024) and a significant strain difference for 70 dB SPL clicks (*p* = 0.0440). Overall, the ABR Wave I data for clicks in quiet and in noise suggest that α9α10KO mice had more robust responses compared to WTs.

To further evaluate the effects of masking noise on suprathreshold α9α10KO ABRs, we derived a ratio comparing Wave I amplitude in quiet and in noise for a given stimulus, level, and subject (*Wave I Amplitude Ratio* = *Amplitude_Quiet_* / *Amplitude_Noise_*). The addition of masking noise is expected to reduce Wave I amplitudes, so amplitude ratio values should be less than 1. Wave I amplitude ratios were similar for WT and α9α10KO mice across most stimulus conditions (Fig 2). Since our masking noise had a constant intensity, the masking noise generally had a greater effect on Wave I amplitudes to lower intensity stimuli (e.g. for the click, Fig 2A).

**Figure 2.**
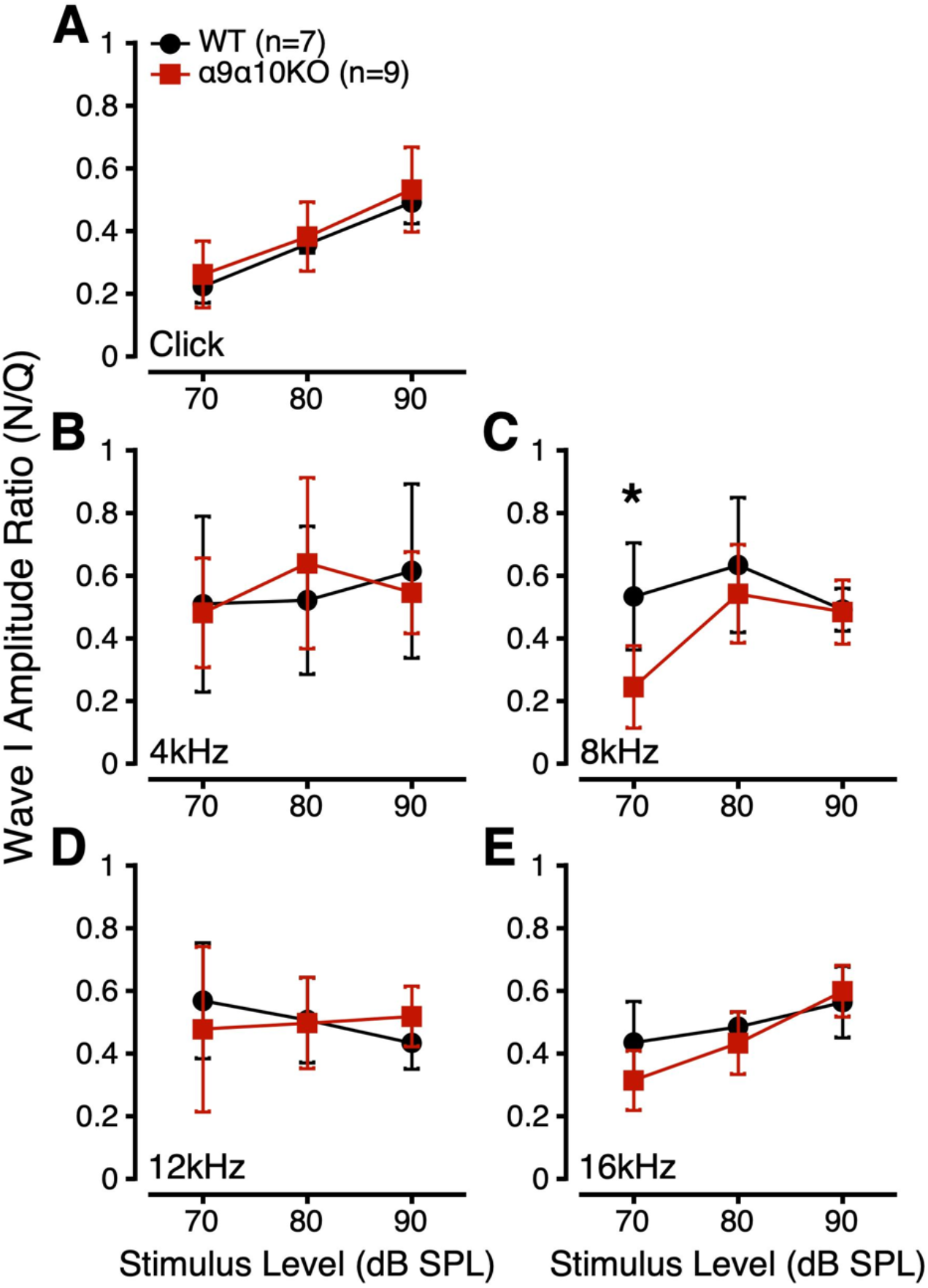
ABR Wave I amplitude ratios (*Amplitude_noise_ / Amplitude_quiet_*) to suprathreshold clicks (A) and tonebursts (B: 4kHz, C: 8kHz, D: 12kHz, E: 16kHz) for α9α10KO (squares) and WT mice (circles).

Linear mixed-effects models were used to compare Wave I amplitude ratios for each stimulus across strain, stimulus level, and sex. For Wave I amplitude ratios to clicks, this analysis revealed a significant main effect of click level and no significant interactions (see Table I). Post hoc analyses indicated significant differences in Wave I amplitude ratios for 70, 80, and 90 dB SPL clicks (*p* < 0.0001 for all). For 8kHz tonebursts, there were significant main effects of strain, sex, and stimulus level, and a significant interaction between strain/stimulus level. Interestingly, post hoc analyses revealed a significant strain difference at 70 dB SPL (*p* = 0.0008; Fig 2C). For 16kHz tonebursts, there was a significant main effect of stimulus level and a significant interaction between strain/stimulus level. Post hoc analyses revealed no significant strain differences at any stimulus level, though there was a trend towards a significant strain difference at 70 dB SPL (*p* = 0.0857). LMEs for 4 and 12 kHz toneburst Wave I amplitude ratios revealed no significant main effects or interactions. Overall, the Wave I amplitude ratio data suggested that suprathreshold α9α10KO ABRs were more susceptible to masking noise at 8kHz than WTs.

### B. Acoustic startle responses in quiet and in noise

ASRs were measured to noise bursts of varying intensity in quiet and in 60 dB SPL masking noise. Figure 3 shows mean ASR amplitudes for each strain as a function of pulse level in quiet (A) and noise (B). ASR amplitudes varied extensively across individual animals of both strains, as is common for mice. As expected, ASR amplitude increased with pulse level across masking conditions and strains. ASR amplitudes were similar for WT and α9α10KO mice in both quiet and in noise. For both strains, the addition of masking noise increased ASR amplitudes for high intensity pulses, consistent with previous reports (Carlson & Willott, 2001; Ison, 2001; Kim et al., 2022; McGuire et al., 2015).

**Figure 3.**
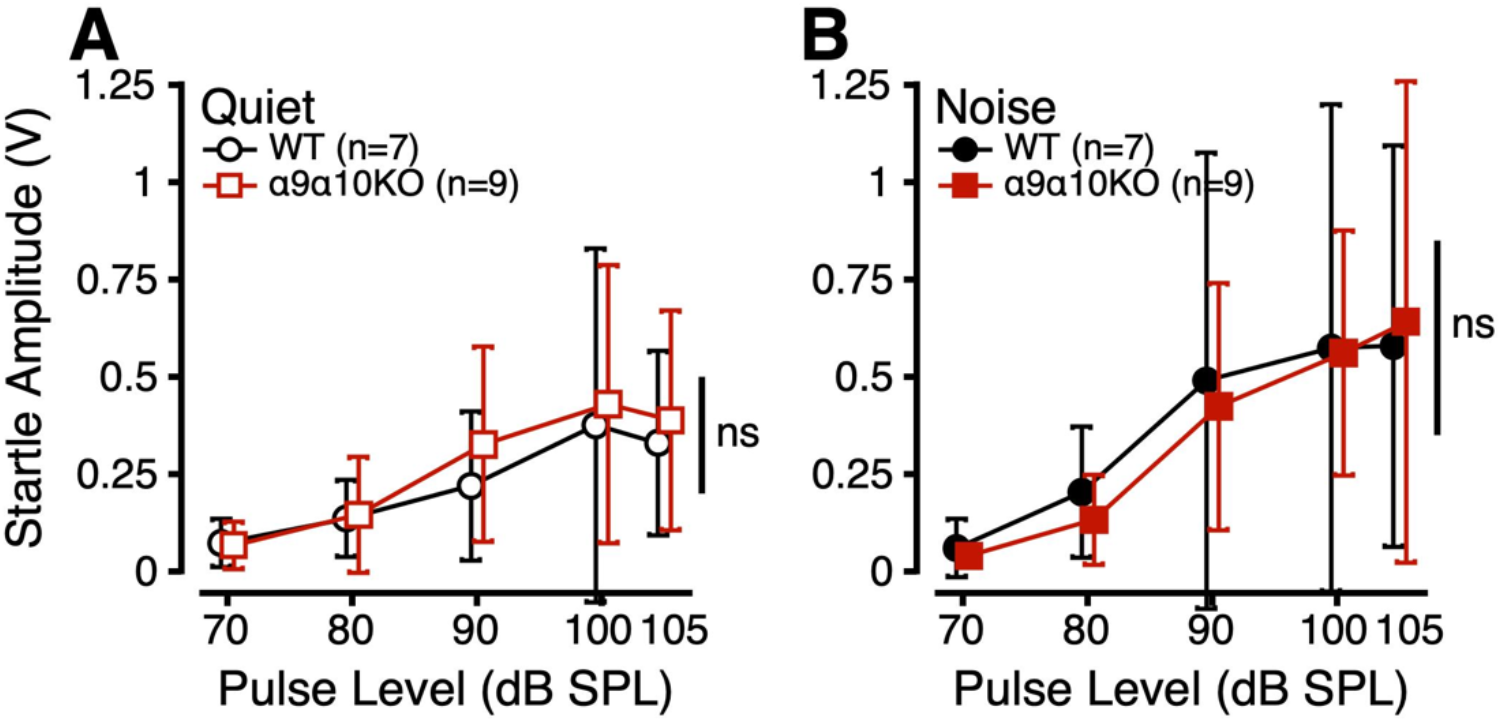
Acoustic startle response amplitudes (V) as a function of pulse level (dB SPL) in quiet (open; A) and in noise (filled; B) for α9α10KO (squares) and WT mice (circles).

Linear mixed-effects models comparing ASR amplitudes in quiet across strain, sex, and pulse level revealed a significant main effect of pulse level (see Table I). Post hoc analyses revealed significant differences in ASR amplitude between pulse levels of 70/90, 70/100, 70/105, 80/90, 80/100, 80/105, 90/100 (*p* < 0.0001 for all), and 90/105 (*p* = 0.239). Similarly, linear mixed-effects models comparing ASR amplitudes in noise revealed a significant main effect of pulse level, with post hoc analyses showing significant differences in ASR amplitude between pulse levels of 70/80 (*p* = 0.0433), 70/90, 70/100, 70/105, 70/105, 80/90, 80/100, 80/105 (*p* < 0.0001 for all), 90/100 (*p* = 0.0464), and 90/105 (*p* = 0.0100). These results suggest that ASR amplitudes peaked at 100 and 105 dB SPL and that there were no differences in ASR amplitudes between WT and α9α10KO mice in quiet or in noise.

### C. Prepulse inhibition of the acoustic startle response: Frequency difference sensitivity

PPI of the ASR was used to probe sensitivity to frequency differences (FD). In this task, startle-inducing noise bursts presented in a 70 dB SPL 10kHz tone background were preceded by a change in background tone *frequency*. If the mouse is able to perceive this prepulse cue, they should inhibit their startle to the oncoming noise burst. Startle amplitudes are plotted as a function of prepulse frequency in Figure 4A. Startle amplitudes were variable across and within strains, but generally decreased as the prepulse frequency deviated farther from 10kHz. α9α10KO mice had larger startle amplitudes across all FD conditions compared to WTs. Linear mixed-effects models comparing startle amplitudes as a function of strain, sex, and prepulse frequency revealed significant main effects of strain and frequency. Post hoc analyses revealed a trend toward a significant difference in startle amplitude across strain (*p* = 0.0538) and significant differences in startle amplitude between prepulse frequencies of 7/10 (*p* = 0.0001), 7/10.5 (*p* = 0.0049), 8/10 (*p* = 0.0046), 10/13 (*p* = 0.0175). These results suggest that α9α10KO mice may be hyperreactive to certain acoustic startle conditions compared to wildtypes.

**Figure 4.**
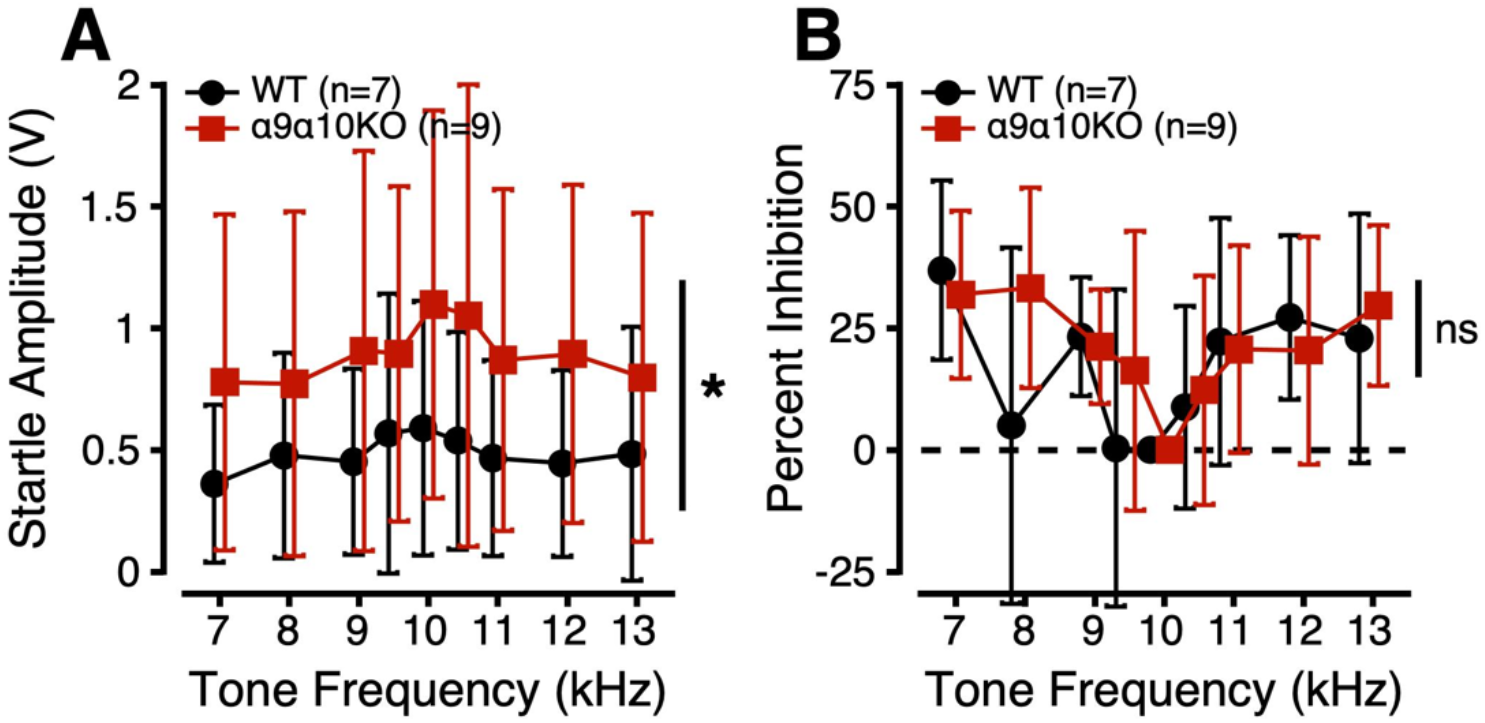
Frequency difference sensitivity estimated from prepulse inhibition of the acoustic startle response for α9α10KO (squares) and WT mice (circles). A. Acoustic startle response amplitudes (V) as a function of prepulse cue frequency (kHz). B. Percent inhibition of the ASR as a function of prepulse cue frequency.

PPI of the ASR was calculated according to: *Percent Inhibition* = (*Startle_no_change_* - *Startle_change_* / *Startle_no_change_*) * 100. When an animal inhibits its ASR, the PPI value should be greater than 0. In Figure 4B, PPI is plotted as a function of prepulse frequency. Despite the α9α10KO mice having larger raw ASR values in the FD task, they showed similar PPI to the WTs (Fig 4B). Linear mixed-effects models comparing PPI as a function of strain, sex, and prepulse frequency revealed a significant main effect of frequency (see Table I). Post hoc analyses revealed significant differences in PPI between prepulse frequencies of 7/9.5 (*p* = 0.0022), 7/10 (*p* < 0.0001), 7/10.5 (*p* = 0.0076), 9/10 (*p* = 0.0174), 10/11 (*p* = 0.0256), 10/12 (*p* = 0.0076), and 10/13 kHz (*p* = 0.0019). These results suggest that WT and α9α10KO mice have similar frequency difference sensitivity.

### D. Prepulse inhibition of the acoustic startle response: Intensity difference sensitivity

PPI of the ASR was also used to probe sensitivity to intensity differences (ID). In this task, startle-inducing noise bursts presented in a 70 dB SPL 10kHz tone background were preceded by a change in background tone *level*. Startle amplitudes are plotted as a function of prepulse level in Figure 5A. As with the FD task, startle amplitudes in the ID task were variable across and within strains, but generally decreased as the prepulse level deviated farther from 70 dB SPL. α9α10KO mice had larger startle amplitudes than WT mice across all ID conditions. Linear mixed-effects models comparing startle amplitudes as a function of strain, sex, and prepulse level revealed significant main effects of strain, sex, and level, but no significant interactions (see Table I). Post hoc analyses revealed significant differences in startle amplitude across strain (*p* = 0.0292) and between prepulse levels of 70/80 (*p* = 0.0347) and 72/80 (*p* = 0.0498). Consistent with the FD testing, these results again suggest that α9α10KO mice may have exaggerated startle responses to some acoustic stimuli compared to WTs.

**Figure 5.**
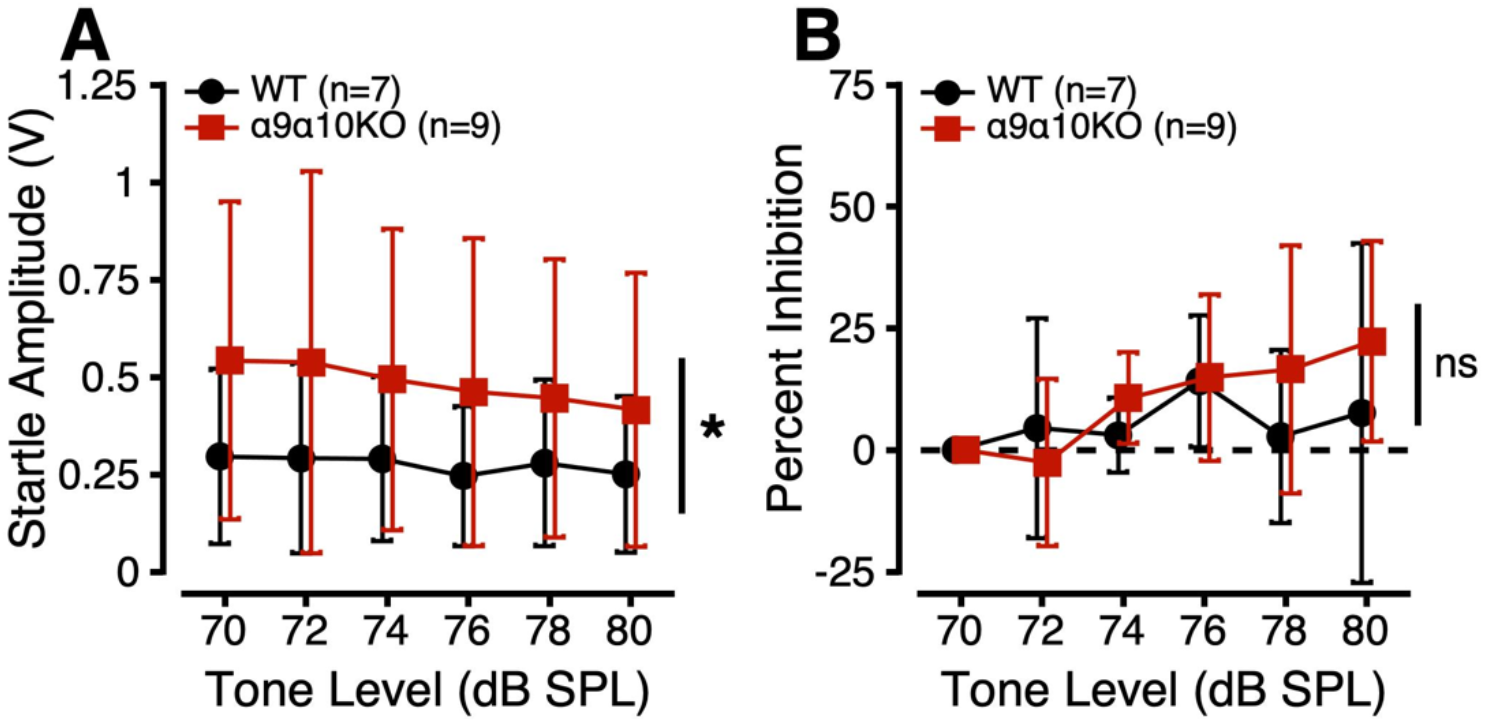
Intensity difference sensitivity estimated from prepulse inhibition of the acoustic startle response for α9α10KO (squares) and WT mice (circles). A. Acoustic startle response amplitudes (V) as a function of prepulse cue level (dB SPL). B. Percent inhibition of the ASR as a function of prepulse cue level.

In Figure 5B, PPI is plotted as a function of prepulse level. Despite the α9α10KO mice having larger raw ASR values in the ID task, they showed similar PPI to the WTs (Fig 5B). Linear mixed-effects models comparing PPI as a function of strain, sex, and prepulse level revealed a significant main effect of prepulse level. However, no post hoc comparisons were significant. These results suggest that WT and α9α10KO mice have similar intensity difference sensitivity.

## IV. DISCUSSION

We further characterized auditory function in the α9α10KO mouse model with deficient medial olivocochlear inhibition of outer hair cells by measuring ABRs in quiet and in noise, ASRs in quiet and in noise, and PPI measures of frequency and intensity difference sensitivity. Consistent with previous findings in α9KOs, α9α10KO mice showed normal ABR thresholds in quiet, normal intensity difference sensitivity, and hyperreactivity to sounds under some ASR and PPI stimulus conditions (Clause et al., 2017; Lauer, 2017; Lauer & May, 2011; Vetter et al., 1999). However, in contrast to predictions about impaired medial olivocochlear function, α9α10KO mice showed normal ABR thresholds in noise, enhanced ABR amplitudes in quiet and in noise, and grossly normal frequency difference sensitivity compared to wildtypes. We infer that α9α10KO mice may develop compensatory mechanisms that support auditory function in the absence of normal MOC function, as has been suggested for α9KO mice (May et al., 2002). This study contributes to the growing, but conflicting literature on the role of the MOC system in hearing in noise (Lauer et al., 2022).

### A. Physiological responses to signals in quiet and noise

We used ABRs in quiet and in noise as physiological estimates of hearing sensitivity and hearing-in-noise function. ABR thresholds in quiet and in noise suggested comparable hearing sensitivity for α9α10KOs and WTs, consistent with previous reports in α9KOs (Lauer, 2017; Lauer & May, 2011; Morley et al., 2017). ABR wave amplitudes and latencies were also examined to evaluate whether morphological differences were apparent across strains. Wave morphology was highly variable, especially for central waves, so we focused our analyses on ABR Wave I only. The larger Wave I amplitudes observed for α9α10KO mice than WTs may be due to the lack of efferent suppression of transient tone stimuli or developmental compensatory processes in the α9α10KOs. Furthermore, a smaller suprathreshold Wave I amplitude ratio (noise/quiet at 8kHz) for the mutants suggested that α9α10KOs were more susceptible to masking noise than WTs, consistent with weaker MOC-mediated noise suppression. Sex effects may have also contributed to the physiological differences between α9α10KOs and WTs (Dondzillo et al., 2021), as male mice exhibited a greater number of strain differences than females. Since C57BL/6J mice lose their MOC responses as early as 8 weeks, despite intact responses at 6 weeks (Zhu et al., 2007), it is possible that strain differences in the ABR would be more apparent in slightly younger animals (Morley et al., 2017).

### B. Behavioral responses to signals in quiet and noise

We measured behavioral responses to sounds in quiet and in noise using ASR and PPI techniques to probe hearing-in-noise function. We utilized these measures to avoid repeated exposure to sounds and practice effects that occur with traditional rodent psychoacoustic tasks out of a concern for potential behavioral compensation (Lauer et al., 2017; May et al., 2002). Previous experiments in α9KO mice on a range of background strains have shown abnormally large prepulse facilitation in response to short gaps in noise (Lauer & May, 2011), reduced PPI in quiet but not noise (Allen & Luebke, 2017), and reduced PPI to changes in frequency, but not intensity (Clause et al., 2017). In contrast, we found no differences in PPI to frequency or intensity changes in α9α10KO mice. Although PPI methodology is limited in its ability to assess acuity (Lauer et al., 2017), we were surprised to find no gross differences here.

However, α9α10KO mice showed larger than normal ASR amplitudes in the presence of constant background tones, indicating increased salience of the startle-eliciting stimuli under these conditions. Animal weights were similar across strains, as were the ASR amplitudes in quiet and broadband noise backgrounds. Thus, we cannot attribute the effect to differences in mass reactivity on the ASR apparatus. The hyperreactivity to abrupt, loud sounds could indicate a form of loudness hyperacusis experienced in the presence of background sounds with starkly different spectral content. Prior studies have shown ASR potentiation in the presence of background sounds (Basavaraj & Yan, 2012; Carlson & Willott, 2001; Gerrard & Ison, 1990; Hoffman & Fleshler, 1963). The background tone might normally trigger efferent suppression of the response to the broadband startle-eliciting stimulus – an effect which is absent in the α9α10KO mice.

### C. Potential compensatory mechanisms in MOC mutant mice

Overall, α9α10KO mice exhibited variable responses to signals in noise, so it is unclear whether they are more or less susceptible to masking effects. The larger ABR Wave I amplitudes and larger ASR amplitudes suggest compensation and/or hyperreactivity in α9α10KO mice. One potential mechanism for preserving hearing function in α9α10KO mice is activity-dependent plasticity in the lateral olivocochlear efferents (Frank et al., 2022; Niu & Canlon, 2002; Wu et al., 2020). A previous study reported qualitatively normal ChAT staining of lateral olivocochlear synapses in α9α10KO mice (Morley et al., 2017; not quantified), but changes in other neurotransmitter systems have not been investigated. It is feasible that, amidst diminished medial olivocochlear feedback, neurotransmitter expression in the lateral efferent neurons adjusts to compensate for the lack of medial efferent mediated cochlear gain control. This could result in preserved ABR thresholds in quiet and noise via direct modulation of auditory nerve activity. The role of the lateral olivocochlear neurons on more complex aspects of hearing behavior is entirely unknown, since specific manipulations of this pathway have not been applied in psychoacoustic experiments (but see Allen and Luebke (2017) for some potential lateral efferent mediated effects involving calcitonin gene-related peptide (CGRP)).

Other compensation mechanisms contributing to α9α10KO phenotypes may include plasticity of the central auditory pathway. The absence of MOC gain control in α9α10KO mice is like having a chronic noise exposure, since background noises will not be attenuated. This also results in an increase in afferent activity. Enhanced auditory input is known to broadly and significantly impact physiology in the brainstem and auditory cortex (Ngodup et al., 2015; Occelli et al., 2022; Oliver et al., 2011; Willott et al., 2005; Zhang et al., 2001). Specific contributions from peripheral and central compensatory mechanisms to hearing-in-noise phenotypes in MOC mutants are unknown and should be explored in future studies.

## V. SUMMARY AND CONCLUSIONS

The persistent difficulty in identifying clear and consistent hearing deficits in genetically mutated mice with impaired medial olivocochlear activity, as well as in behaviorally trained animals with surgical lesions of the medial olivocochlear system (e.g. Dewson, 1967; Igarashi et al., 1972; May & McQuone, 1995; Trahiotis & Elliott, 1970), underscores the need for more specific, controlled, acute manipulations to elucidate the effects on hearing. It is perhaps unsurprising that different functional outcomes have been reported in α9KO, α10KO, and α9α10KO strains, given differences in genetic background, development, age at testing, environmental housing conditions, ambient sound exposure histories, and test measurements across studies. Genetically engineered strains (like α9α10KOs mutants) inherently develop with diminished or enhanced medial olivocochlear activity. To overcome this limitation, future experiments could make use of virally introduced genetic gain-of-function mutations (Zhang et al., 2023), inducible gene knockouts, DREADDS, or optogenetic stimulation or silencing of medial olivocochlear neurons in adult animals. Additionally, potential compensatory mechanisms such as lateral olivocochlear plasticity should be further investigated, as these mechanisms have important implications for listeners with degenerating medial olivocochlear systems due to age, noise, and other damaging circumstances (Lauer, 2017; Lauer et al., 2022; Vicencio-Jimenez et al., 2021).

## VI. ACKNOWLEDGMENTS

The authors would like to acknowledge that this research was supported by NIH NIDCD R01 DC017620 (PI: Lauer), NIH T32 DC000023 (Mondul; PIs: Cullen, Fridman), NIH NIDCD F32 DC020346 (PI: Burke), the Nebraska Tobacco Settlement Biomedical Research Foundation (Morley), and the David M. Rubenstein Fund for Hearing Research (Lauer). The cloning and initial generation of the mouse strains were conducted by Genoway, Inc. (Lyon, France).

## VII. AUTHOR DECLARATIONS

The authors do not have any conflicts of interest to disclose. All procedures were approved by the Johns Hopkins University Animal Care and Use Committee (ACUC) and follow the NIH ARRIVE Guidelines.

## VIII. DATA AVAILABILITY

The data that support the findings of this study will be publicly available on the Johns Hopkins University data repository following publication of the manuscript. Interested parties may contact the authors for additional data inquiries.

